# Distribution of native European spiders follows the prey attraction pattern of introduced carnivorous pitcher plants

**DOI:** 10.1101/410399

**Authors:** Axel Zander, Marie-Amélie Girardet, Louis-Félix Bersier

## Abstract

Carnivorous plants and spiders are known to compete for resources. In North America, spiders of the genus *Agelenopsis* are known to build funnel-webs, using *Sarracenia purpurea* pitchers as a base, retreat and storage room. They also very likely profit from the insect attraction of *S. purpurea*. In a fen in Europe, *S. purpurea* was introduced ~65 years ago and co-occurs with native insect predators. Despite the absence of common evolutionary history, we observed native funnels-spiders (genus *Agelena*) building funnel webs on top of *S. purpurea* in similar ways as *Agelelopsis.* Furthermore, we observed specimen of the raft-spider (*Dolmedes fimbriatus*) and the pygmy**-**shrew (*Sorex minutus*) stealing prey-items out of the pitchers. We conducted an observational study, comparing plots with and without *S. purpurea*, to test if *Agelena* were attracted by *S. purpurea*, and found that their presence indeed increases *Agelena* abundance. Additionally, we tested if this facilitation was due to the structure provided for building webs or enhanced prey availability. Since the number of webs matched the temporal pattern of insect attraction by the plant, we conclude that the gain in food is likely the key factor for web installation. Our results provide an interesting case of facilitation by an introduced plant for a local predator, which has developed in a very short time scale.

## Introduction

Introduced species, especially predators, are known to have many negative effects on local fauna (e.g. Gurevitch and Padilla 2004; Salo et al. 2007; Paolucci et al. 2013; Zander et al. 2015). Studies reporting positive effects (e.g. facilitation or mutualism) of an introduced species for one or more native species are rarer (e.g. Bially and Macisaac 2000; Castilla et al. 2004; Müller-Schärer et al. 2014). In her review, Rodriguez (2006) identified several pathways by which novel species can facilitate native species. Among these are novel habitat creation (e.g. Schwindt et al. 2001) and trophic subsidy, like nutrient enrichment (e.g. Quinos et al. 1998), increase of prey availability (e.g. Ortega et al. 2004), or the possible usage of new host plants (e.g. Graves and Shapiro 2003). Furthermore the release of competition and predation (e.g. Knapp et al. 2001; Grosholz 2005) play an important role for facilitation of native species by introduced ones. The replacement of existing species is especially important when considering the potential value of alien species for conservation (e.g. Angradi et al.; Schlaepfer et al. 2011). In case the species that is replaced by the introduced species has become rare and only the new species can provide the facilities, like nesting places (e.g. Zavaleta et al. 2001) or simply shadow and shelter (Van Riel et al. 2000), some other native species will depend on the introduced species in order to survive.

In Europe, despite their differing phylogenetic backgrounds, the shrew *Sorex minutus*, the funnel web spiders of the genus *Agelena* (either *Agelena labyrinthica or Agelena gracilens* (*syn.: Allagelena gracilens*)), and the raft spider *Dolomedes fimbriatus* have similar food spectra, preying mostly on insects (Pernetta 1976; Nyffeler and Benz 1978; Poppe and Holl 1995). While *Agelena sp.* and *Sorex minutus* are found in many habitats, *Dolomedes fimbriatus* is specialized in semiaquatic habitats, e.g. bogs and fens (Nyffeler and Benz 1978). *Sarracenia purpurea* is a carnivorous plant species originating from North America, and has been introduced to European bogs and fens (Zander et al. 2015). This could create direct competition for prey, but also the possibility for predation facilitation, for example through kleptoparasitism by the above mentioned species.

Prey abundance and availability of web building sites are factors limiting growth and survival of spiders (e.g., Colebourn 1974; Wise 1979; Whitney et al. 2014; Takada and Miyashita 2014). Competition for web building sites is common and well described (Riechert, 1978, 1981; Wenseleers et al. 2013; Opatovsky et al. 2016). Locally higher prey abundance is known to affect individual spider fitness (Wise 1975, 1979), and may cause spiders to build their webs closer to each other. However a minimal distance will be maintained, probably to reduce the exploitative competition between spider-individuals. In Europe, an average minimum distance between webs of ca. 60 cm was observed for *Agelena labyrinthica* (Fasola and Mogavero 1995).

Prey availability can be increased by the presence of carnivorous plants (Wolfe 1981; Zamora 1995). Furthermore spiders can be facilitated by the provided structure to build their webs (Milne 2012). However carnivorous plants and spiders can also act as antagonistic resource competitors i.e. through kleptoparasitism (robbing of already caught prey from the plant; (Zamora 1995; Jennings et al. 2010). Cresswell (1993) gives a further example for resource parasitism between linyphiid spiders and the carnivorous pitcher plant *Sarracenia purpurea.* Here the spiders build their webs directly over the pitcher openings, as such preventing insect to fall in the traps. Furthermore in North America, the native range of the *S. purpurea* plants, funnel-webs spiders of the genus *Agelenopsis* are known to build their webs in the way that the funnels end in the pitchers of *S. purpurea,* not only blocking the entrance for possible prey, but also using pitchers as a base, storage room and retreat (Milne 2012). Both linyphiid and *Agelenopsis* spiders profit very likely from the insect attraction of *S. purpurea* and are thus facilitated by the plant.

Prey attraction by *S*. *purpurea* is a locally and also a temporally limited event. *S*. *purpurea* produces nectar at the opening of the pitcher leaves to attract insects (Deppe et al. 2000; Bennett and Ellison 2009). Nectar production starts early after pitcher opening, and ceases a few weeks later. Consequently, pitchers catch most of their prey in the first few weeks after opening (Fish and Hall 1978; Wolfe 1981; Heard 1998). Capture rates raise very fast after pitcher opening and peak after 3-4 weeks, then decrease also quite fast to reach a low level plateau around 35-40 days after opening (Fish and Hall 1978; Heard 1998).

The funnel-web spider genus *Agelena* has a similar ecology than the American *Agelenopsis*. Despite the short evolutionary history between *Agelena* and *S. purpurea,* we observed webs built on the plant in Champ Buet. *Agelena-*spiders are generalist predators (Tanaka 1991). For example, the diet of *Agelena labyrinthica* consists of insects from 12 orders, mainly Orthopthera, Coleoptera, Hymenoptera, Lepidoptera and Diptera (Nyffeler and Benz 1978). *Agelena* species build their funnel-shaped webs close to the ground and are thus catching both flying and walking insects (Nyffeler and Benz 1978). For this reason, compared to orb-web spiders, their diet is likely more similar to that of *S. purpurea*, which consists mainly of Diptera, Hymenoptera, Coleoptera, Hemiptera, and Aranea (Judd 1959; Cresswell 1991; Heard 1998; Newell and Nastase 1998).

Here, we explore the possible interactions between *S. purpurea* and *Agelena* spiders. We conducted a study in the Champ Buet site, where we compared six 25 m^2^ plots with and without *S. purpurea* plants. We first analyse if the presence of *S. purpurea* increases the local abundance of *Agelena* spiders. Furthermore, we asked whether spiders use *S. purpurea* because they offer suitable structure and shelter, or because of the increased prey availability provided by insect attraction. If the latter proposition is correct, we expect the abundance of *Agelena* spiders to match the temporal pattern of insect attraction of *S. purpurea*. Specifically, we tested the following hypotheses: 1) at the local scale, *Agelena* spiders are more abundant in plots with *S. purpurea*; 2) spider numbers change proportionally to insect attraction patterns of *S. purpurea*; 3) during insect attraction period, spider webs are not randomly distributed within *Sarracenia* plots, but are aggregated close to or on the *S*. *purpurea* plants.

## Materials and Methods

### Study site

All field experiments were done in Champ Buet (46°36′46.35″N / 6°34′42.75″E ca. 600m above sea level, see Fig. Sb 4b). We obtained authorisation from the Canton of Vaud to conduct research. The site, which is partly dominated by *Eriophorum latifolium* and partly by *Phragmites australis*, is mowed once a year to prevent shrub encroachment. Around 65 years ago, *S. purpurea* was introduced to Champ Buet from Les Tenasses (ca. 1250 m), where the population was already well established and in a growing state (Parisod et al. 2005). Although the pitchers tips are mowed off once a year, plants cope quite well with the local conditions.

### Study system

*Sarracenia purpurea* was introduced in Europe and Switzerland during the 19^th^ Century. *S. purpurea* was planted in various bogs and fens located in the Jura mountains and in the Alps in the late nineteenth century (Correvon 1947). *S. purpurea* is a carnivorous pitcher plant, naturally occurring in swamps and peat bogs in North America. Often, plants reproduce vegetatively and form clumps of several individuals. In Champ Buet, the largest clumps reached a diameter of approximately 1 m. To provide some extra nutrients, the leaves of *S. purpurea* are pitcher shaped and serve as traps mostly for arthropods. To attract prey, *S. purpurea* secrete an extra- floral nectar in the upper part of the pitchers (Deppe et al. 2000; Bennett and Ellison 2009). This secretion stops after some weeks. Thus, after this time, pitchers become less attractive for potential prey organisms. Capture rates raise very fast after pitcher opening, peak after 3-4 weeks, then decrease also quite rapidly to reach a low level around 35-40 days after opening (Fish and Hall 1978; Heard 1998, see inset of Fig. Sa 1-5).

The funnel-web spiders in the present study were determined by Gilles Blandenier to be juveniles of the Genus *Agelena*. In Switzerland, two *Agelena* species occur, the more common *Aglena labyrinthica* and *Agelena gracilens* (syn. *Allagelena gracilens*) (Info Fauna 2016), which cannot be distinguished in the juvenile stage (Blandenier personal communication). The study was performed in early summer, thus only juveniles occur (Fasola and Mogavero 1995). Both species build a flat net that extends into a funnel shaped tube (Fig. 1 and Fig. Sb 1-3). In this tube, the spider hides and waits for prey.

**Fig. 1:**
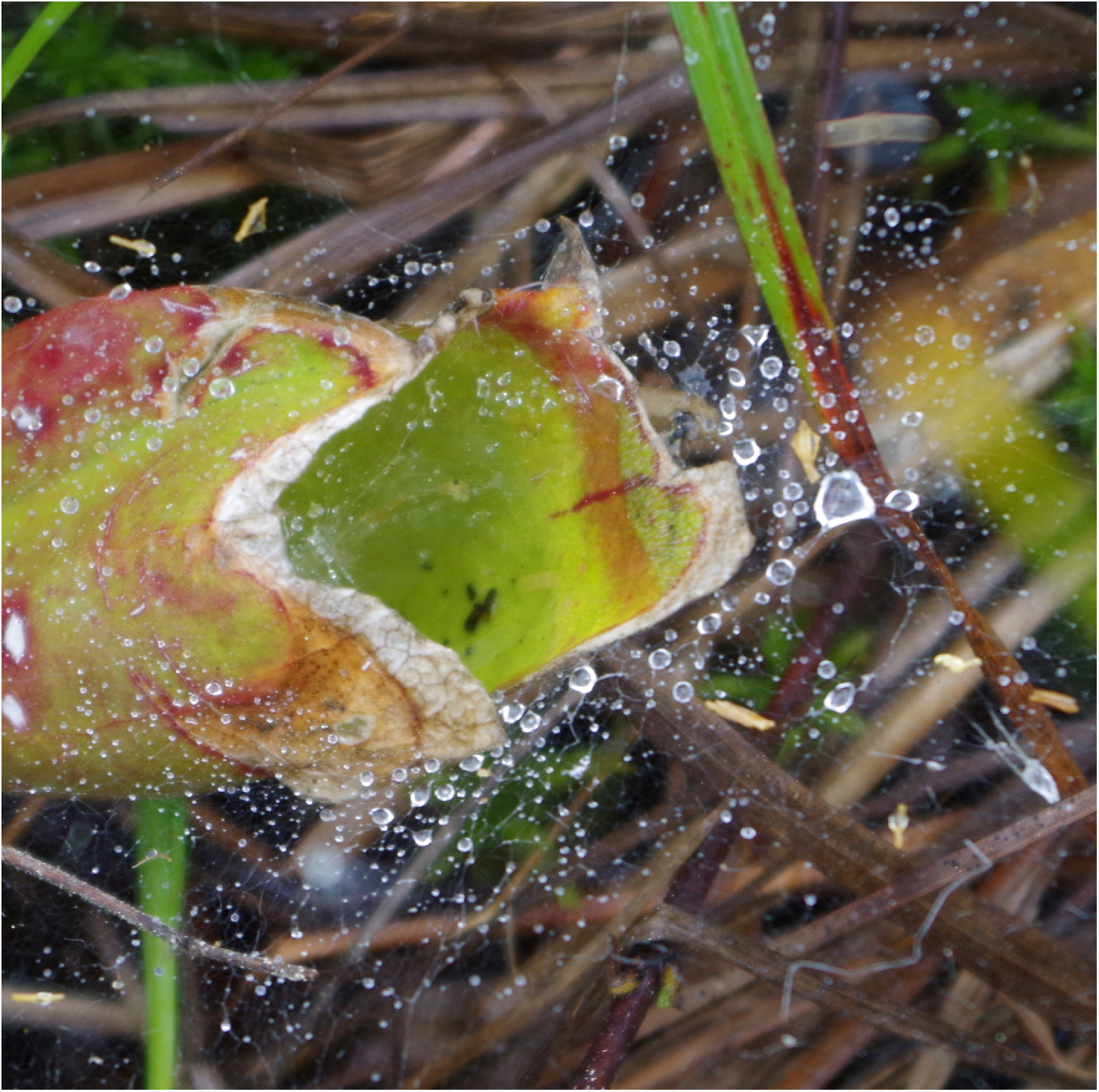
*Agelena* sp. web funnelling directly into a *S. purpurea* pitcher (tip mowed off).

Since both species have similar ecology in terms of web construction and diet, we consider individuals at the genus level in our study.

Two other species interacting with *S. purpurea* occurred in our sites, for which we report our observations in the Result section. The raft spider *Dolomedes fimbriatus* (Pisauridae) is a big semi-aquatic spider that is found near many types of freshwater habitats, like streams, lakes and fens (Carico 1973). Although these spiders feed occasionally on aquatic vertebrates and invertebrates (Bleckmann and Lotz 1987), the main prey of *Dolomedes fimbriatus* are terrestrial invertebrates (Poppe and Holl 1995). Females can reach up to 23mm in body length and are thus able to catch quite big prey items, like dragonflies. *Sorex minutus*, the smallest mammal of central Europe, is a quite common and widely distributed insectivore, mainly feeding on various arthropods and mollusks (Pernetta 1976). In the Swiss Alps, it inhabits a wide array of habitats (wet meadows, marshes and mixed forests) as well as altitudinal belts up to 2496m (Marchesi et al. 2014). We observed it in the site of Les Tenasses (ca. 1250m *46°29′28.51″N, 6°55′16.04″E)*, where *S. purpurea* has become invasive in the ombrotrophic zone of the bog (Feldmeyer 1985).

### Experimental set-up

We randomly chose six 5×5 m plots, with the constraint that three contained *S. purpurea* plants, three had no *S. purpurea* and, to prevent interference between plots, that the minimum distance between them was 10 m. Plots were marked with a plastic stripe that did not affect arthropod movements. All plots were further divided in twenty-five 1 m^2^ quadrants (subplots). The presence/absence of *S. purpurea* plant was recorded for each subplot (see Fig. Sa 1-5), and the number of clumps inside each 25 m^2^ plot was counted. Subplots without *S. purpurea* were additionally separated in two classes: subplots neighbouring a subplot with *S. purpurea* (“close”), or subplots further away (“away”).

When the majority of the pitchers began to open, we recorded the numbers and distribution in each subplot of all *Agelena* webs and followed the changes over time. Sampling occurred at least biweekly, with some variability, from the 19^th^ of May till 14^th^ of July (5 sampling sessions). Note that webs of other spider species were present, but their number was low and negligible compared to *Agelena* webs, and they were not considered.

### Measured variables

At each sampling session, we determined the total number of webs in each 1 m^2^ subplot and in the 25 m^2^ plots, accordingly. These abundances were used as response variables. We also noted if webs were built directly on clumps of *Sarracenia* or not (substrate). The attribute of the plots (with or without *S. purpurea*) and of the subplots (with or without *S. purpurea*; close and away), and the substrate for the webs were used as explanatory variables.

Additionally, we collected prey in pitchers and determined the specimens under dissecting microscope at the highest possible taxonomic level. Sampling was performed by removing all material from 35 randomly selected pitchers of the same age, at days 14, 22, and 36 after opening.

### Statistical analyses

When analysing the relationships between the dependent variable “number of webs” (log-transformed) and 1) the types of experimental plots (with or without *Sarracenia*), 2) the type of substrate (on *Sarracenia* plants or on other structures), and 3) the number of *Sarracenia* clumps (only for plots with *Sarracenia*). To account for repeated measures (5 sampling sessions), we used a first-order auto-regressive (AR1) autocorrelation structure in the residuals (Zuur et al. 2009). All model residuals were checked for normality with QQ plots. Analyses were performed with the gls function of the nlme package (Pinheiro et al. 2011) in R (R Core Team 2013).

When comparing observed and expected frequencies of webs built 1) in subplots close to vs. away from subplots with *Sarracenia* for all sampling sessions, and 2) on Sarracenia vs. not on Sarracenia for each sampling session, we used log-likelihood ratio tests with Williams correction (Sokal and Rohlf 1995). We applied a sequential Bonferroni correction of p-values for the latter test (Holm 1979).

## Results

### Number of Agelena webs in plots with and without Sarracenia

The presence of *Sarracenia* attracted more *Agelena* spiders than are usually found in 25m^2^ plots without *Sarracenia* in the Champ Buet field side. Over all sampling sessions, there were significantly more webs present in plots with *Sarracenia* compared to those without (Fig. 2, Table Sa 1). The numbers of webs inside the Sarracenia plots increased with time (except for the last sampling day; see Fig. 3), and peaked at 3-4 weeks after leaf opening, which corresponds to the time point when pitchers catch the most insects (Fish and Hall 1978; Heard 1998, see Fig. Sa 1 to 5, and Fig. Sa 6). In contrast, the number of webs in the plots without *Sarracenia* slightly decreased (Fig. 3). This interaction over time of webs in plots with Sarracenia and webs in plots without was significant (Bonferroni corrected p-value = 0.026; see also Fig. Sa 7).

**Fig. 2:**
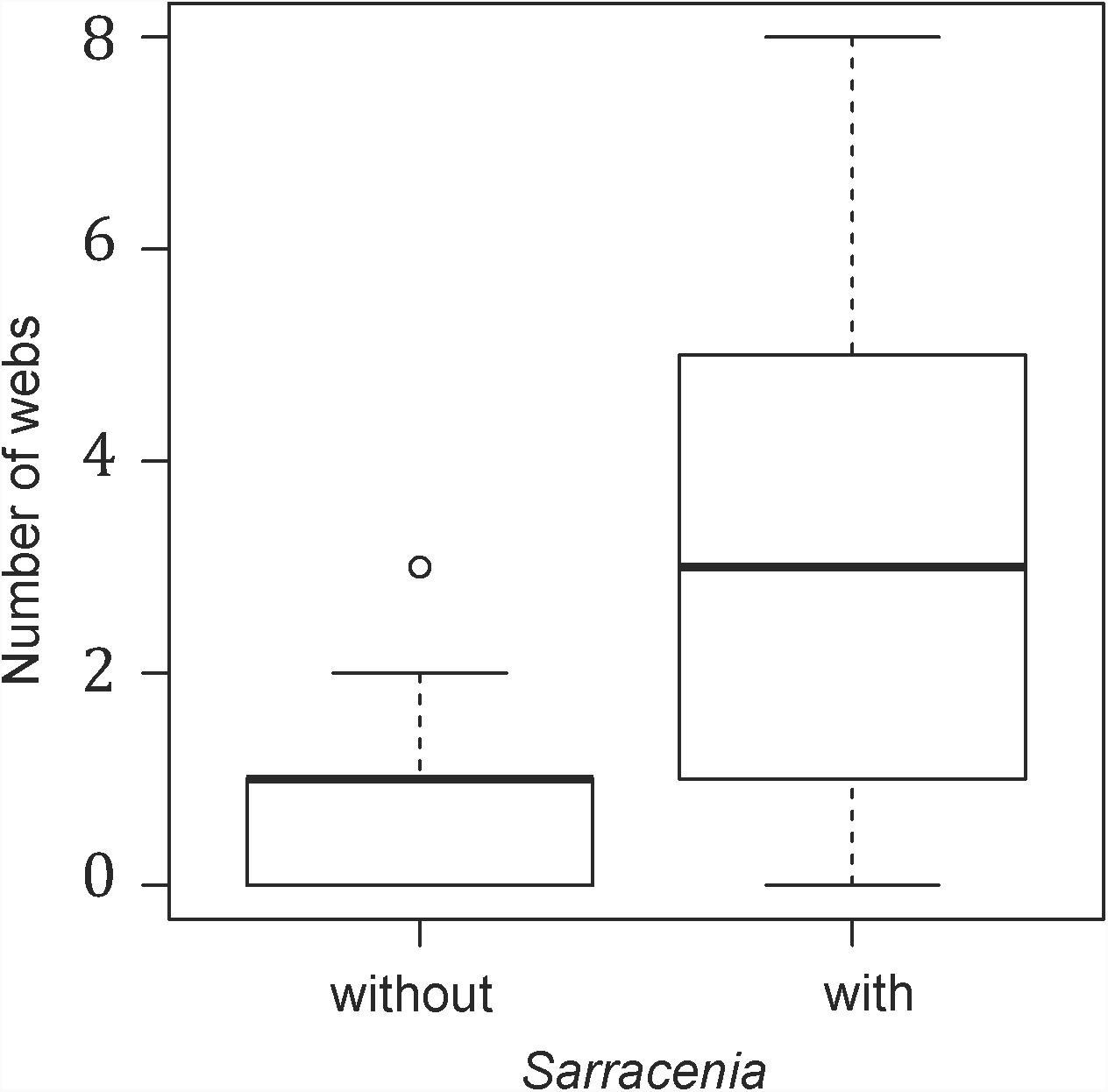
Comparison of numbers of webs in plots with and without *Sarracenia*. In *Sarracenia* plots there were significantly more webs than in the plots without *Sarracenia* (generalized least squares model with AR1 correlation structure for repeated measures; p-value < 0.001, see Table S1a for details).

**Fig. 3:**
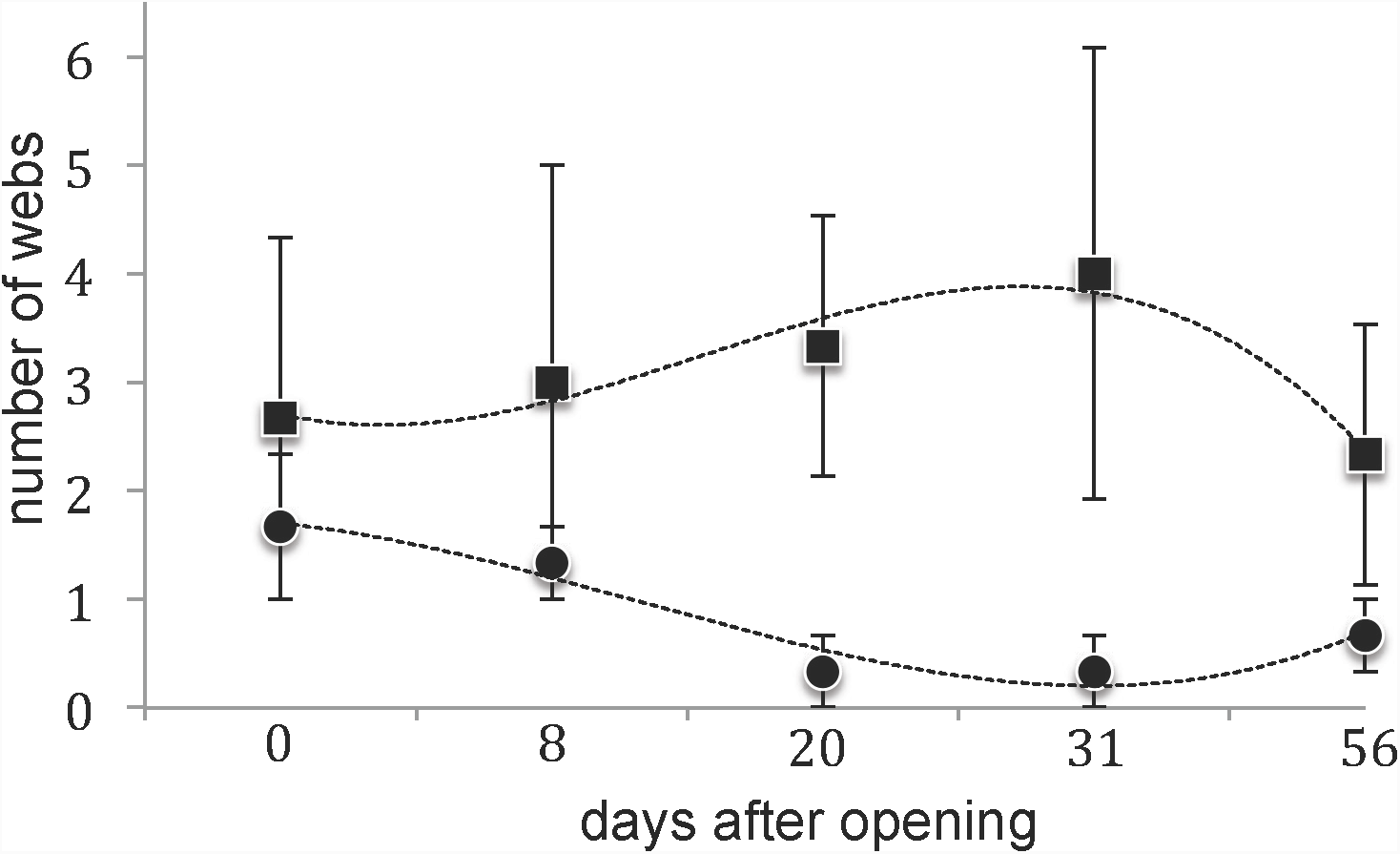
Average number (with standard error) of webs in 25 m^2^ plots with *Sarracenia* (squares) and without *Sarracenia* (circles) over time (days after average date of pitcher opening). During the peak of insect attraction (day 20-31) the differences in plots with and without *Sarracenia* were highest. Dotted lines (third order polynomial regressions) are added to guide the eye. The numbers of webs in both types of plots (a and b) are negatively correlated (Spearman rank correlation (for data without “day 56”); rho = -0.95, p-value = 0.05).

### Number and distribution of Agelena webs in plots with Sarracenia

Inside the *Sarracenia* plots, the abundance and distribution of *Agelena* webs were not random. First, according with the findings of Fasola and Mogavero (1995) the minimum distance between two webs of *Agelena* was always bigger than 59 cm, resulting in the fact that no *S. purpurea* clump was occupied by the webs of two spiders at a time.

Second, the number of webs were positively correlated with the number of *Sarracenia* clumps inside a plot (generalized least-square model with AR1 correlation structure for repeated measures; p-value < 0.001, see Fig. Sa 8,). Third, *Agelena* spiders build their webs either directly in 1m^2^ subplots with *Sarracenia* plants, or in neighboring subplots (see Fig. Sa 1-5). Consequently, there were no webs in subplots more than one meter away from a *Sarracenia* clump (Fig. 4; likelihood-ratio test for the expected number of webs in subplots neighbouring *Sarracenia* clumps vs. in subplots at least one meter away from *Sarracenia* clumps; p-value = 0.004; see Table Sa 3).

**Fig. 4:**
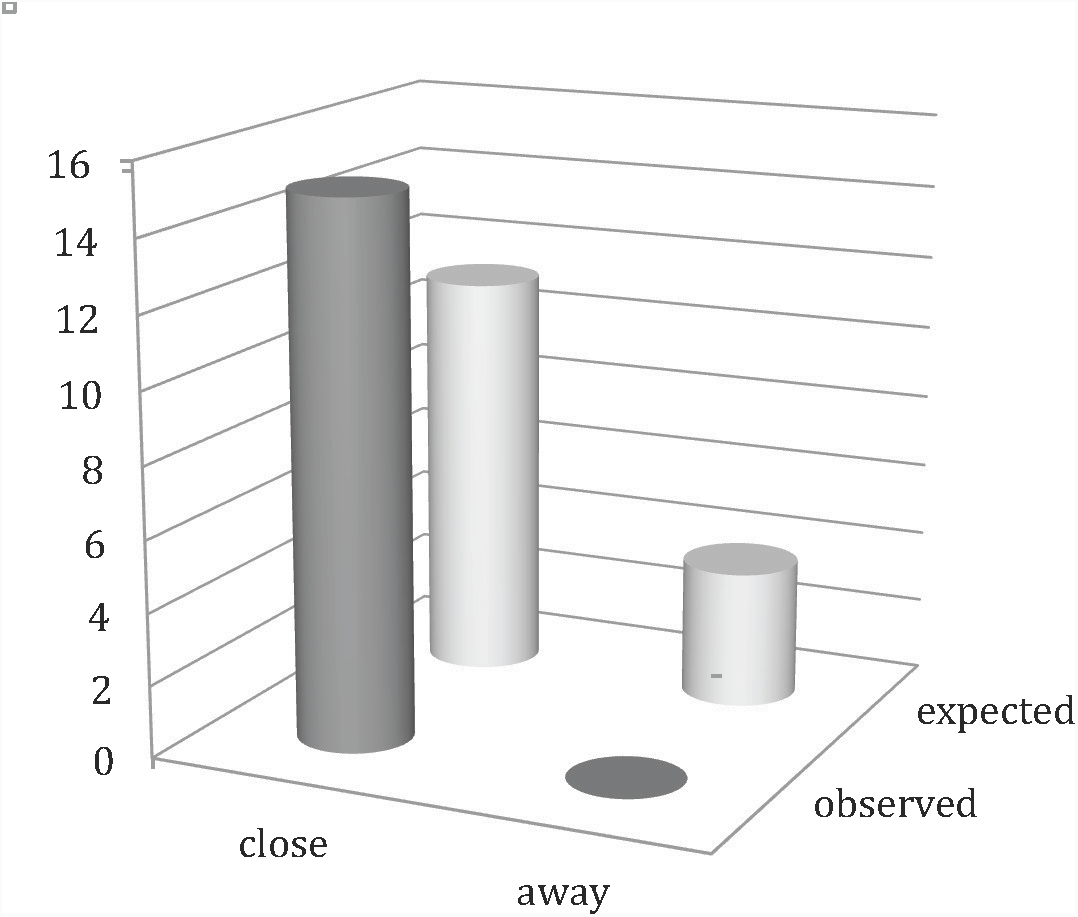
Contrary to expectations of random distribution, webs in subplots without *Sarracenia* are always located close to subplots with *Sarracenia* (consequently, there were no webs in subplots more than one meter away from a *Sarracenia* clump; likelihood-ratio test, p-value = 0.004, see Table S3 for details).

Fourth, when considering all measured time points, we recorded a significantly higher number of cobwebs built directly on *Sarracenia* clumps compared to the rest of the plot (Fig. Sa 9). This effect is particularly strong given that the actual number of subplots covered with *Sarracenia* plants is much smaller than the number that is not (global ratio: 1 : 3.7; see Table Sa 4 for results of generalized least square model).

### Number and distribution of Agelena webs over time

At start of the experiment the ratio of webs directly on *Sarracenia* compared to the webs in the rest of the plots was 50:50. However, during the peak of insect attraction (sampling sessions 3 and 4), over 80% of the webs were found on the plants. This number fell again to 70% after insect-attraction decreased (Fig. 5a). Except for day 0, all days showed a significantly higher proportion of webs directly on *Sarracenia* (Bonferroni corrected p-values: day 0, 0.42; day 8, 0.022; day 20, < 0.001; day 31, < 0.001; day 56, 0.031; see also Table Sa 5). Even when the actual number of webs is considered without correcting for the availability of space for building webs (Fig. 5b), there were from day 8 on more webs on *Sarracenia* than in the rest of the plot. This effect became less strong at the end of the experiment, when insect attraction by *Sarracenia* is decreased (Fig. Sa 6; see also Fish and Hall 1978; Heard 1998).

**Fig. 5a:**
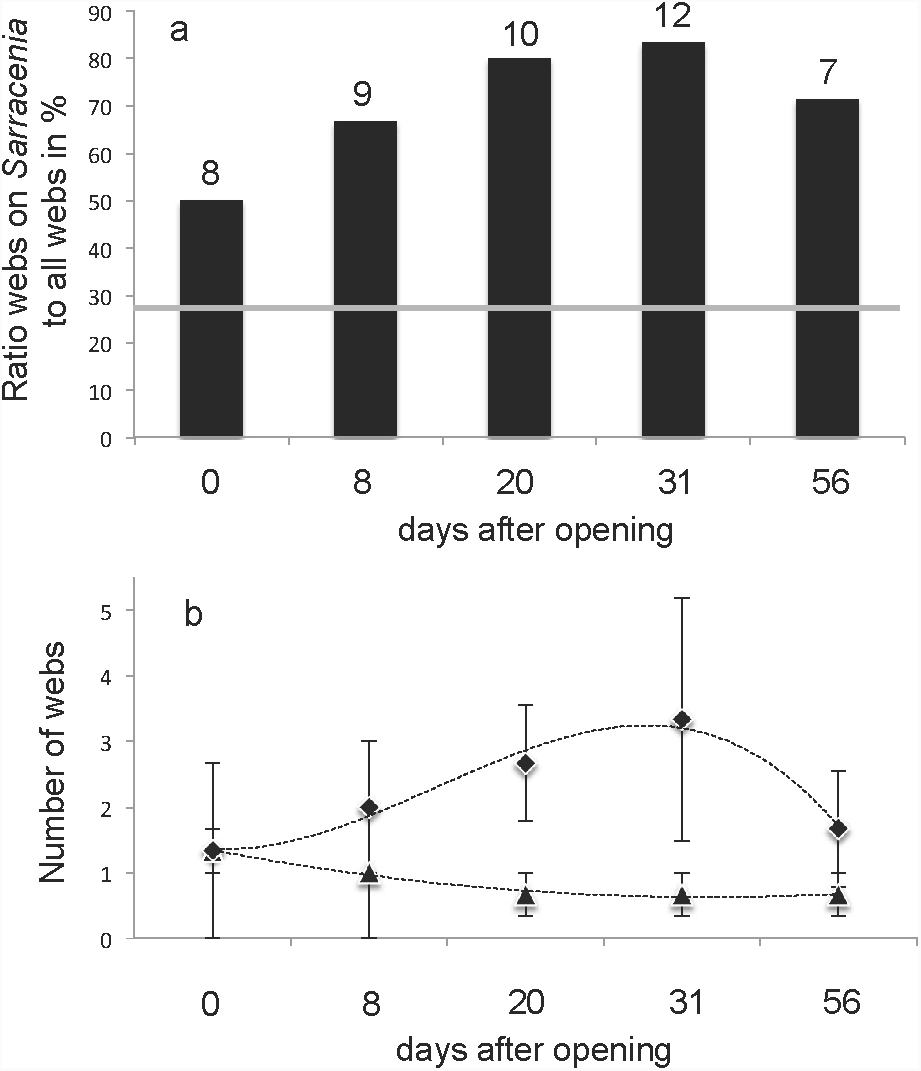
Ratio of webs directly on *Sarracenia* plants over webs not on *Sarracenia* (measured for plots with Sarracenia only), followed over time since the estimated date of leaf opening. Total number of webs in the plots at that day is given on top of the bars. The line shows the expected ratio assuming a random distribution of the webs. Likelihood ratio tests for each sampling day showed significantly more webs in Sarracenia subplots, except for day 0 (sequential Bonferroni corrected p-values: day 0, 0.42; day 8, 0.022; day 20, < 0.001; day 31, < 0.001; day 56, 0.031; see Table S 5 for details on statistics). **b:** Average number of webs on *Sarracenia* plants (diamond) compared to average number of webs elsewhere in the plots (triangles), over time (average and standard error). We observe an increase for webs on *Sarracenia*, except for the last sampling date, and a steady decrease for webs not on *Sarracenia*. Dotted lines (third order polynomial regressions) are added to guide the eye.

### Kleptoparasitism by Dolomedes and Sorex

*Dolomedes fimbriatus* were found regularly luring next or even inside *S. purpurea* pitchers for prey (see Fig. Sb 5-9). Adult *Dolomedes* spiders can climb in and out of the pitchers with ease, even when the pitchers are water filled. We observed some spiders removing big insect prey items from the pitchers. We observed only one adult *Dolomedes* spider drowned in a pitcher (see Fig. Sb 10), and one juvenile individual (Supplemental Material c), although their nests were sometimes located surrounded by *S. purpurea* clumps (Fig. Sb 11).

In the Site of Les Tenasses, the shrew *Sorex minutus* was observed diving headfirst into *S. purpurea* pitchers and feeding on prey items directly in the pitcher (Zander, personal observation).

### Prey of Sarracenia

During our prey survey in 2012 we found 229 prey items in the 35 sampled pitchers of the Champ Buet field site. More than 95% of them were insects (See Fig. 6 and Supplemental Material c, Table S6); only 6 spiders from the families Salticidae, Pimoidae (*Pimoa*), Thomisidae (*Xysticus*) and Lycosidae (*Pardosa*), 2 mites and one mollusc were found in the pitchers. Note that no *Agelena* spider was found as prey of *S. purpurea.* The prey-numbers caught over time follow closely the distribution described by Heard (1998) (Fig. Sa 6, see also Fig. Sa 1-5). We collected 39 prey items two weeks after pitcher opening, 114 after three weeks, and 76 after five weeks. Diptera were by far the most abundant prey order (135 individuals; see also Fig. 6) followed by Coleptera and Hymenoptera.

**Fig. 6:**
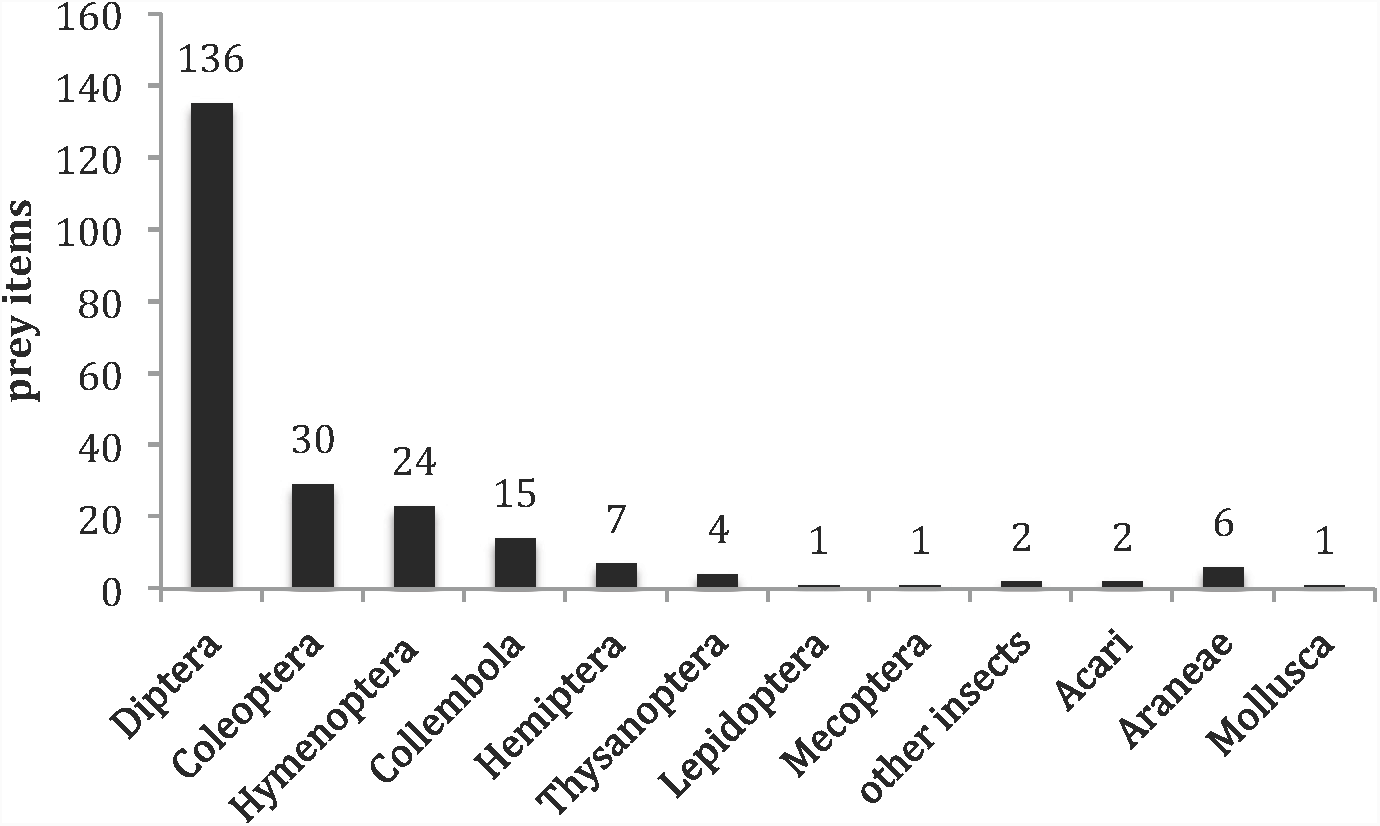
*S. purpurea* prey of 35 pitchers in Champ Buet for the first 6 weeks after opening. Insects from the order Diptera are by far the most abundant prey items.

These result are consistent with studies from the natural range (Judd 1959; Cresswell 1991; Heard 1998; Newell and Nastase 1998; Wallen 2008) and show that insect-prey attraction mechanisms work worldwide. Also, for Europe, our data match the observations of Hartmeyer (1996), Adlassnig et al. (2010) and Fragnière (2012). Note that although the 35 pitchers had the same age, we observed a high variability in prey capture rates: 9 pitchers caught no prey, 15 up to ten, and the maximum was 26 prey items.

## Discussion

While the functional role of *S. purpurea* is that of a predator (Fig. 6; also see Supplemental Material c), it acts more as a facilitator than as a competitor for other predatory species. Our results indicate that individuals of *Agelena* are facilitated by the *S. purpurea* presence in terms of habitat modification (creating structure for building webs) and increased prey availability. It is commonly observed that introduced species can be beneficial to some types of local organisms while harming another ones. For example, the invasive zebra mussel *Dreissena polymorpha* is detrimental to several native species, but is also facilitating local arthropods through habitat creation by forming dense colonies, and acting as refuge (Bially and Macisaac 2000). Refuge creation by *S. purpurea* might also be important for *Dolomedes fimbriatus.* Furthermore, during the limited period of insect attraction by the plant’s nectar production (Heard 1998, Deppe et al. 2000, Bennett and Ellison 2009), *Agelena, Dolomedes* and *Sorex* are facilitated by increased prey availability (trophic subsidy, see Rodriguez 2006). Since these species were never or rarely (2 specimens of *Dolomedes*) found as prey in pitchers, our results confirm the facilitative role of *S. purpurea* for *Agelena* spiders, and to a lower extent for *Dolomedes* spiders.

In the *Agelena* experiment, we found the facilitation effect to act at different spatial scales. More webs were found in plots with *S. purpurea*; inside plots with *S. purpurea*, we observe a positive relationship between plant and spider abundances; finally, webs were not randomly distributed, but placed either directly on or in close vicinity of the plants. This positive effect on *Agelena* densities did not change the territorial behaviour of the spiders, and we never found two webs on the same *S. purpurea* clump (see also Fasola and Mogavero 1995). Although *S. purpurea* facilitates *Agelena* spiders in two ways, firstly by providing suitable structure for web building, and secondly by increasing prey availability, our results show that the latter factor is of key importance. The increased prey availability is due to nectar induced insect attraction of *S. purpurea* (Deppe et al. 2000; see also Fig. Sb 14), which occurs only during a short time frame (Heard 1998). We found the abundance of *Agelena*- webs rising and falling proportionally to the magnitude of the observed insect attraction pattern of *S. purpurea* over time, giving support to the increased prey availability being the main factor for the facilitative effect. Our results also provide indication of a global rearrangement of spider web distribution in the field site driven by *S. purpurea* pitcher phenology. *Agelena* spiders are attracted from the surrounding environment towards the *S. purpurea* clumps when insect attraction levels are high, and disperse back to the bog after insect attraction ceased.

To our knowledge this is the first time that this behaviour of *Agelena* was shown in Europe. Note that we found no other sites in Switzerland where funnel spiders co-occur with *S. purpurea*; this is because the other locations all are at higher altitudes, where climatic conditions are unfavourable for *Agelena* spiders, which occur only in warm habitats. Most interestingly however, the genus *Agelena* occurs only in the Old World, while the genus of *Sarracenia* originates from North America. The very similar spider genus of *Agelenopsis* occurs in North America, and these spiders were already observed building funnel-webs that use *S. purpurea* pitchers as retreat and even storage room for prey (Milne 2012). Yet, these American species coexisted at least since the last glaciation, while the interaction of *Agelena* sp. with *S. purpurea*, including the described behavioural change in the hunting activities, can only have had developed during the last 65 years in Champ Buet. From the literature, other spider-pitcher interactions are known. For example, in South East Asia, the spider *Misumenopsis nepenthicola* is known to capture prey inside the pitcher leaves of *Nepenthes gracilis* (Chua and Lim 2012), a behaviour similar to our observations of *Dolomedes fimbriatus*. Common garden experiments with *Agelena* and *Dolemedes* individuals from different sites could reveal if these behaviours are local adaptions.

Our prey survey indicates that *S. purpurea* has a negative impact on the local arthropod community in the bog of Champ Buet. When we extrapolate the number of prey items caught by 35 pitchers to the total amount of pitchers in the site, as a gross estimate 7000 prey items are captured within six weeks. While the main strategy of insect attraction is by nectar production, we observed some pitchers to develop a fetid smell due to a large amount of decaying insects. This lead to the conspicuous attraction of a large number of necrotroph insects like Calliphoridae sp. and *Necrophorus vespilloides* (Adlassnig et al. 2010). However, if too numerous, the decaying prey can even be detrimental to the plant (Adlassnig et al. 2010): we also observed some pitcher leaves full of insects decaying with their prey (personal observation).

Our study describes a remarkable case of facilitation by an introduced plant for a local predator. This result raises interesting ecological questions about the possible benefits for *S. purpurea*. For example, spiders may protect the plant against herbivore attacks from orthopterans or even Stylommatophora (Bruggisser et al. 2012), or prevent overfilling of the leaves and their subsequent negative effects. Also, as mentioned above, the described behaviour between spiders and *S. purpurea* might lead to inherited changes, as is likely the case in the native range of the plant. Finally, such novel interactions with introduced plants may be of conservation importance for endangered species, like *Dolomedes fimbriatus* in our study site. These questions should be examined in further experiments evaluating the fitness consequences of these interactions.

